# Se-Glargine II. Native Function of a Basal Insulin Analog Stabilized by an Internal Diselenide Bridge

**DOI:** 10.1101/2023.06.24.546337

**Authors:** Yen-Shan Chen, Balamurugan Dhayalan, Yanwu Yang, Orit Weil-Ktorza, Norman Metanis, Michael A. Weiss

## Abstract

Insulin glargine, the active component of basal clinical pharmaceutical formulations Lantus® and Toujeo® (Sanofi), provides a model for principles of therapeutic protein design. Formulated in solution at pH 4, insulin glargine exhibits a shifted isoelectric point (from pH 5.3 to neutral pH) due to a basic dipeptide B-chain extension (Arg^B31^-Arg^B32^). In the first article in this series, we described pairwise substitution of Cys^A6^ and Cys^A11^ by seleno-cysteine (Sec; the 21^st^ encoded amino acid) by solid-phase peptide synthesis. ^1^H-^2^H amide proton exchange, as monitored by ^1^H-NMR spectroscopy, provides evidence that substitution of internal cystine A6-A11 by a diselenide bridge stabilizes the protein and damps segmental conformational fluctuations. Further, this analog and its major metabolites M1 and M2 (respectively denoting proteolytic derivatives lacking Arg^B31^-Arg^B32^ or Thr^B30^-Arg^B31^-Arg^B32^) exhibit native hormonal activity in mammalian cell-based assays measuring dose-dependent autophosphorylation of the insulin receptor (pIR/IR ratio) and metabolic gene regulation in human liver-derived HepG2 cells. The internal diselenide bridge also did not alter respective baseline mitogenicities of insulin glargine or its proteolytic products as evaluated by a qPCR-based assay of the balance between proliferative and antiproliferative cyclin gene expression; the assays employed L6 myoblasts over-expressing mitogenic IR isoform A. Given such native function, shelf life—and hence global access to insulin in the developing world—may be enhanced by stabilizing diselenide chemistry.

---

Basal insulin analog formulations are a mainstay of therapeutic regimens in Type 1 diabetes mellitus (T1D; basal-bolus therapy) and in insulin-requiring Type 2 diabetes mellitus (T2D; ordinarily in the absence of rapid-acting formulations). Whereas classical long-acting formulations of animal insulins employed microcrystalline suspensions (such as historic lente insulins and neutral protamine Hagedorn [NPH]; for review, see ref 1), modern basal products contain modified insulin analogs (2). Examples are provided by acylated insulins whose action is protracted by albumin binding and enhanced stability of the subcutaneous (SC) depot (3,4). In the present series of studies, we have focused on insulin glargine, the active component of once-a-day formulations Lantus^®^ and Toujeo^®^ (respective strengths U-100 and U-300; Sanofi). In broad and growing clinical use in the developed and developing worlds (5,6), insulin glargine undergoes pH-dependent precipitation in the SC depot to provide a long-lived depot (7). The analog, formulated in solution at pH 4, contains a dipeptide B-chain extension (Arg^B31^-Arg^B32^) and a substitution at the acid-labile C-terminal residue of the A chain (Asn^A21^→Gly) (Fig. 1).

**Figure 1.**
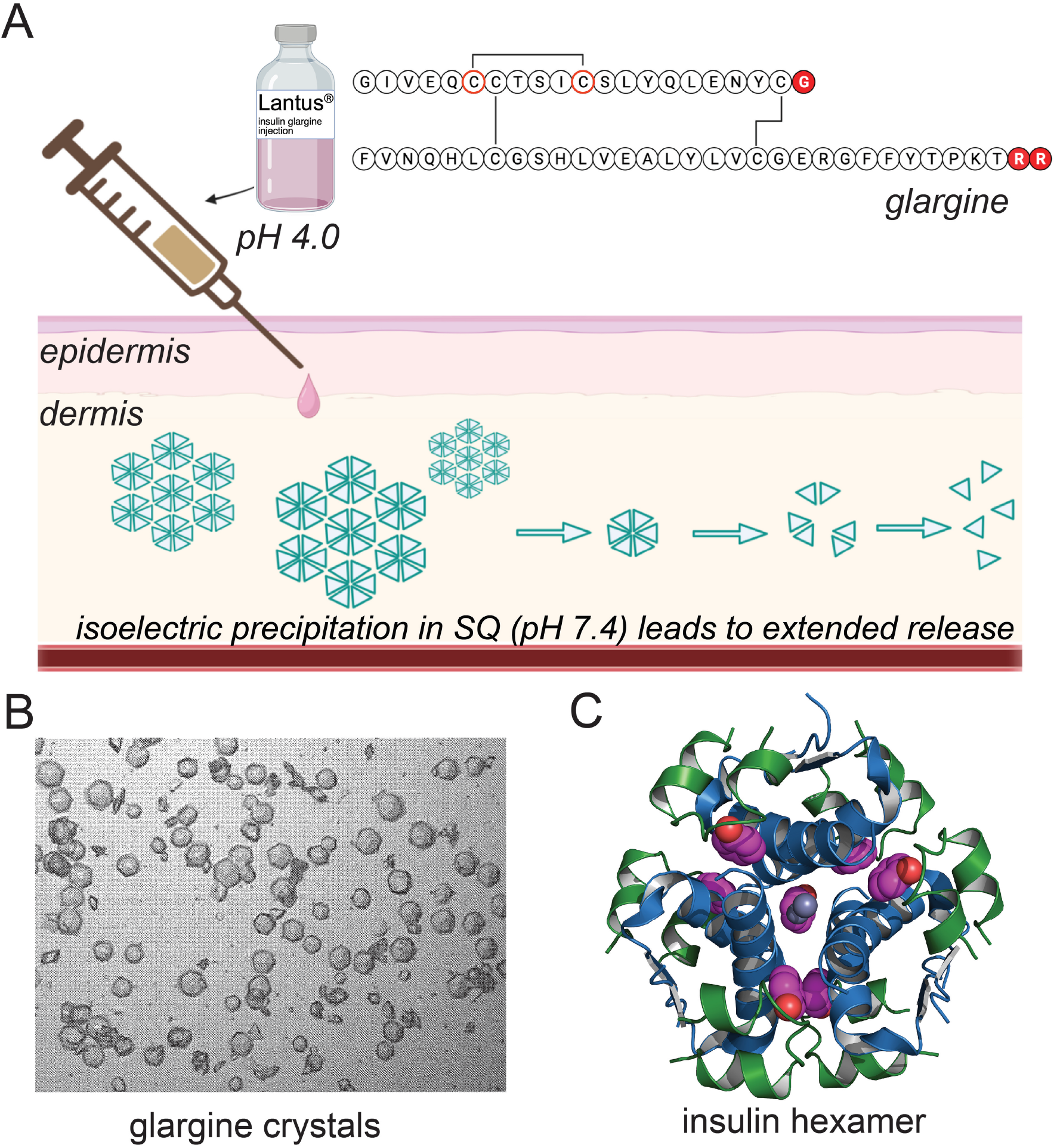
Structure and mechanism of action of insulin glargine. (A) Amino acid sequence and disulfide bond connectivity of insulin glargine (*top right*). Highlighted in red (*filled circle*) are the changes when compared with regular insulin. Cysteine A6 and A11 are highlighted by red circle as the point of selenocysteine modification. Insulin glargine is modified with Arg-Arg tag at the C-terminus of B-chain which shifts its pI close to 7.0. Due to this shift, glargine is formulated at pH 4 to allow its solubility (*top left*). Other modification is due to the deamidation of Asn^A21^ at acidic pH which can be avoided by Gly^A21^ modification. Up on subcutaneous injection, insulin glargine forms microcrystalline precipitates of glargine hexamers (B) which then dissociates slowly to dimers and monomers. (B) hexagonal shaped crystals of glargine in the presence of zinc and phenol (44). (C) A representative crystal structure of insulin hexamer containing zinc and phenol ions (PDB 1ZNJ). A chain (*green*), B chain (*blue*), Zn ions (*blue*) and bound phenol molecules (CPK model, magenta).

In the preceding article (8) we described the chemical synthesis of an analog of insulin glargine in which cystine A6-A11 is substituted by a diselenide bridge (designated “*Se-glargine*”) (9). To this end, Cys^A6^ and Cys^A11^ were pairwise substituted by selenocysteine (Sec), considered the “21^st^ encoded amino acid” (10). The A6-A11 diselenide bridge is compatible with native structure and function of wild-type insulin (9). The present study provides ^1^H-NMR evidence, based on amide-proton exchange in an acidic D_2_O solution, that the modified bridge enhances dynamic and thermodynamic stability. Moreover, the hormonal activities of Se-glargine are essentially indistinguishable as probed in a human liver-derived cell line (HepG2 hepatoma cells; 11). Because insulin glargine is a *prodrug*—the active form in the bloodstream comprises proteolytic products M1 (Gly^A21^-insulin) and M2 (Gly^A21^-*des*-Thr^B30^-insulin)— we also prepared Sec^A6^, Sec^A11^-M1 and Sec^A6^, Sec^A11^-M2. Although unperturbed in activity, M1/M2 derivatives were fortuitously found to be of medical significance given the enhanced mitogenicity (and potential carcinogenicity; 12) conferred by the prodrug’s basic Arg^B31^-Arg^B32^ extension (13). Together, these findings suggest that Se-glargine may be bioequivalent to glargine in the hormonal control of metabolism with advantageous stability properties as a pharmaceutical formulation.

## Results

Glargine insulin modified with selenocysteines at A6 and A11 (Se-glargine) prepared by chain combination as described in the previous paper (see bioRxiv I; ref 8). Briefly, A-chain modified with Selenocysteines at A6 and A11 were prepared by solid phase peptide synthesis on Gly^A21^ resin. B-chain disulfonate was prepared by sulfitolysis of glargine from Lantus^®^ Solostar^®^ pen. Equimolar B-chain disulfonate and [Se^A6^-Se^A11^]-Gly^A21^ A-chain were reacted in pH 11.2 glycine buffer in presence of dithiothreitol. After liquid chromatography/mass spectrometry (LC-MS) indicated formation of product, purification by reversed-phase high-performance liquid chromatography (rp-HPLC) yielded Se-glargine (8).

### Se-glargine exhibits slower ^1^H-^2^H amide exchange

^1^H-NMR spectra of Se-glargine indicated similar, but not identical chemical-shift dispersion with native-like packing of the hydrophobic core. Comparative spectra of native glargine (*top, black*) and Se-glargine (*bottom, red*) are shown in Figure 2A. Although patterns of ^1^H-NMR secondary shifts are similar, spectral line narrowing was observed among (a) upfield-shifted methyl resonances (at right in Fig. 2A) and downfield main-chain amide resonances (at left). Because such resonances in native glargine reflect conformational broadening ascribed to millisecond motions in the protein, we investigated the comparative dynamics and stability of Se-glargine (*versus* native glargine) by ^1^H-^2^H amide-proton exchange in an acidic D_2_O solution (10 mM deuterio-acetic acid at pD 3.0 [direct meter reading] and 25 °C) (14). Resonance assignments were obtained by standard methods.

**Figure 2.**
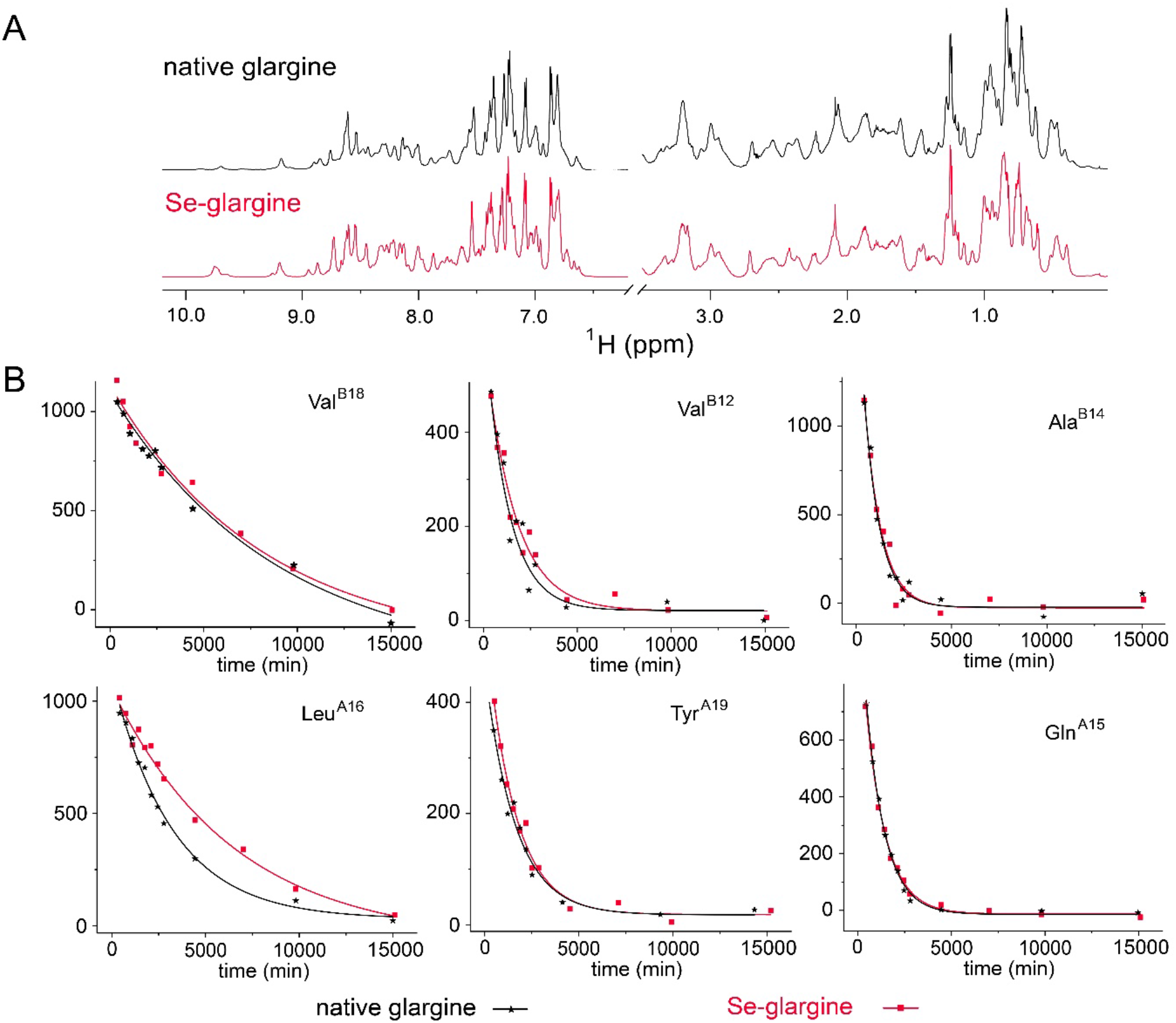
NMR ^1^H-^2^H exchange of native glargine and Se-glargine. (A) 1D ^1^H-NMR spectra of native glargine (*top, black*) and Se glargine (*bottom, red*). Spectral line narrowing was observed in the upfield-shifted methyl and amide/aromatic regions for Se-glargine. Spectra were recorded at a ^1^H frequency of 700 MHz in 10mM deuterated acetic acid (pH 3.0, direct meter reading) at 25 °C. (B) Representative examples of exponential ^1^H-^2^H exchange at specific residues associated with slower exchange kinetics as defined in the native glargine (*black star*) and Se-glargine (*red filled square*). Data were acquired and analyzed by 2D amide-proton ^1^H-^2^H Exchange spectra in 10 mM deuterated acetic acid (pH 3.0, direct meter reading) at 25 °C.

Amide resonance intensities were monitored based on 2D ^1^H_N_-H_α_ cross-peaks in TOCSY spectra. Protection factors (PF) were obtained for six individual amide protons (positions B12, B14, B18, A15, A16, and A19; see Table 1), based on technical accessibility (*i*.*e*. resolved in 2D TOCSY spectra and slowly exchanging). The overall pattern of protection was similar in Se-glargine and native glargine: corresponding sites within the C-terminal A-chain α-helix and central B-chain α-helix as previously observed in studies of insulin *lispro* (14). These respective cross-peak intensives are shown in Figure 2B as a function of time in D_2_O solution. In general, the Se-glargine exhibits slower ^1^H-^2^H exchange. The highest protection factor (PF) belongs to Val^B18^, which adjoins cystine B19-A20 in the hydrophobic core; respective estimates of global stability (ΔG_u_) are 3.66 ± 0.05 kcal/mol (Se-glargine) and 3.46 ± 0.04 kcal/mol (native glargine) (highlighted in red in Table 1). Delayed exchange at the other five sites presumably represents damping of segmental α-helical fluctuations by the diselenide bridge. Such subglobal damping presumably reflects more efficient core packing by the modified bridge (see Discussion).

**Table 1.** ^1^H-NMR Amide-Proton Exchange Studies in D_2_O^a^

| residue | PF | $\Delta G^b$ | PF | $\Delta G^b$ |
| --- | --- | --- | --- | --- |
|  | native glargine |  | Se-glargine |  |
| A16 HA | 232.7 ± 11.1 | 3.23 ± 0.03 | 313.9 ± 14.1 | 3.40 ± 0.03 |
| A19 HA | 169.0 ± 8.7 | 3.04 ± 0.03 | 203.9 ± 19.0 | 3.15 ± 0.05 |
| A15 HA | 98.0 ± 3.1 | 2.71 ± 0.02 | 105.4 ± 3.6 | 2.76 ± 0.02 |
| B18 HA | 346.5 ± 22.4 | 3.46 ± 0.04 | 489.2 ± 38.1 | 3.66 ± 0.05 |
| B14 HA | 129.1 ± 11.5 | 2.88 ± 0.05 | 144.4 ± 18.5 | 2.94 ± 0.08 |
| B12 HA | 60.6 ± 15.2 | 2.43 ± 0.15 | 97.2 ± 11.3 | 2.71 ± 0.07 |

<sup>a</sup><sup>1</sup>H-NMR spectra were acquired in 10 mM deuterio-acetic acid (pD 3.0, direct meter reading) at 25 °C.
<sup>b</sup>Nominal free energies in kcal/mole based on equation $\Delta G = RT \ln(PF)$ . The longest-lived amide resonance in D<sub>2</sub>O with highest protection factor (PF; B18, highlighted in red) is presumed to represent global exchange and so correspond the protein's overall thermodynamic stability as described (14). Comparative values thus provide an estimate of the difference in stability of 0.20(±0.09) kcal/mole ( $\Delta\Delta G_u$ ). The other five sites (black) presumably represent subglobal sites of exchange, providing probes of segmental conformational fluctuations within $\alpha$ -helices.

### Chemical Synthesis of Se-Glargine M1 and M2

Although insulin glargine is itself more mitogenic than wild-type (WT) insulin (12), it is converted into metabolites M1 (*des*-dipeptide[B31-B32]) and M2 (*des*-tripeptide[B30-B32]) in the subcutaneous depot (14). The latter derivatives, the major circulating form of the analog, exhibit mitogenic profiles similar to WT insulin (15).

To assess the corresponding mitogenic properties of Se-glargine, its metabolites were prepared according to *Scheme 1*, wherein an analog of *des*-octapeptide[B23-B30] insulin (DOI) containing the [Se^A6^-Se^A11^]-Gly^A21^ modifications was combined with appropriate B-chain-derived peptides: octapeptide GFFYTOT (for M1) or heptapeptide GFFYTO (for M2). The latter peptides were joined to Arg^B22^ by trypsin-mediated semi-synthesis (16). (In Scheme 1 ornithine (O) was used in place of lysine at position B29 to avoid introducing a second tryptic site (17).) The DOI moiety was obtained from [Se^A6^-Se^A11^]-Gly^A21^-DesDi, a synthetic single-chain insulin precursor (18). After resin cleavage, the 49-residue peptide was folded to form two disulfide bridges (B7-A7 and B19-A20) and one diselenide bridge (A6-A11; Fig. 3). Trypsin cleavage then yielded the 43-residue [Se^A6^-Se^A11^]-Gly^A21^-DOI (Fig. 4) and in turn (under separate conditions) semi-synthetic products (Fig. 5), enabling respective purification of Se-M1 (Fig. 6) and Se-M2 (Fig. 7).

**Scheme 1.**
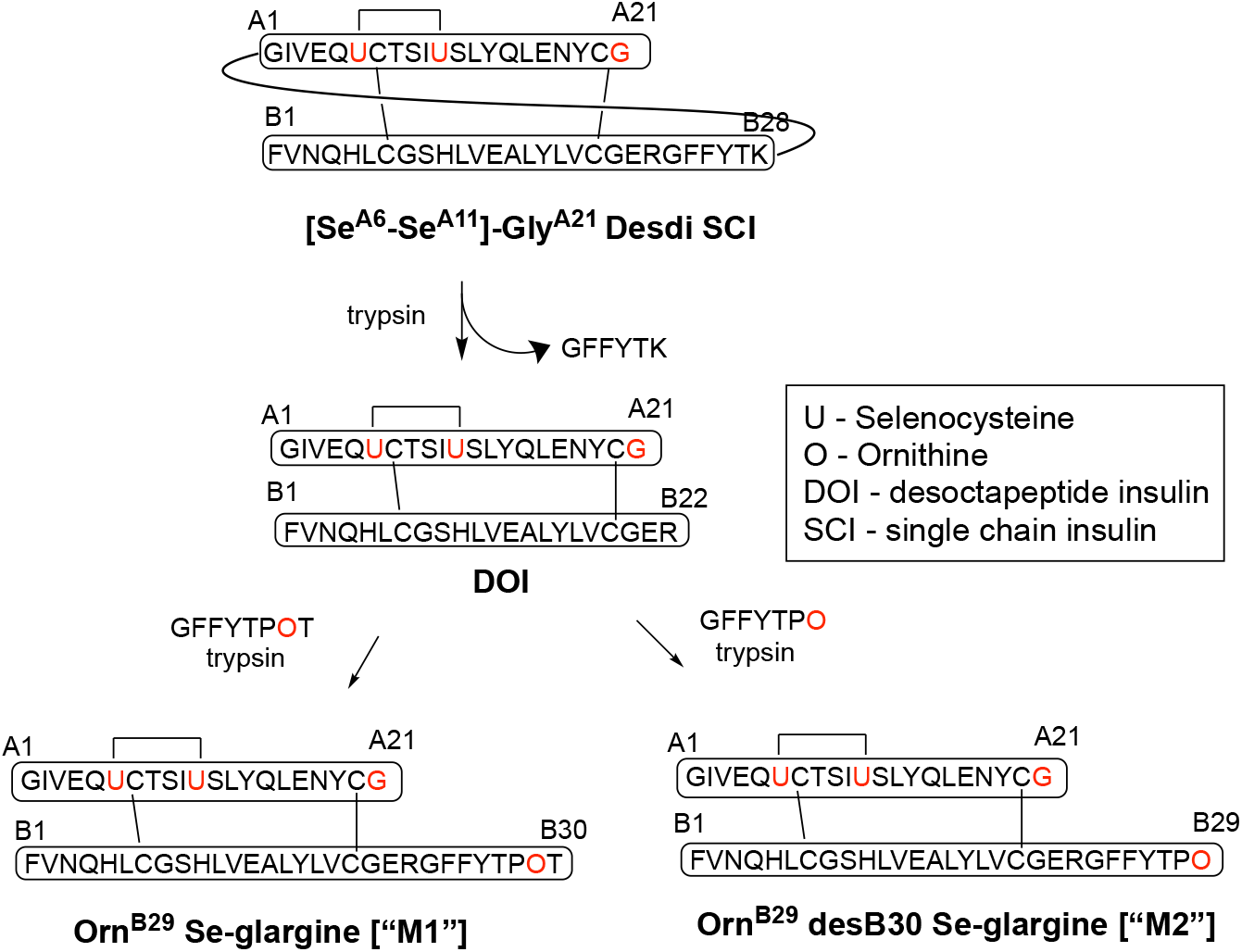
Overview of Se-glargine M1 and M2 synthesis. 1) single chain polypeptide with Gly^A21^ and Se^A6^-Se^A11^ modifications were first folded before subjecting to trypsinization to yield 43-residue desoctapeptide insulin (DOI). 2) The DOI was then used to prepare Se-glargine M1 and M2. B29-Lysine was replaced with Ornithine to avoid cleavage by trypsin during semi-synthesis.

**Figure 3.**
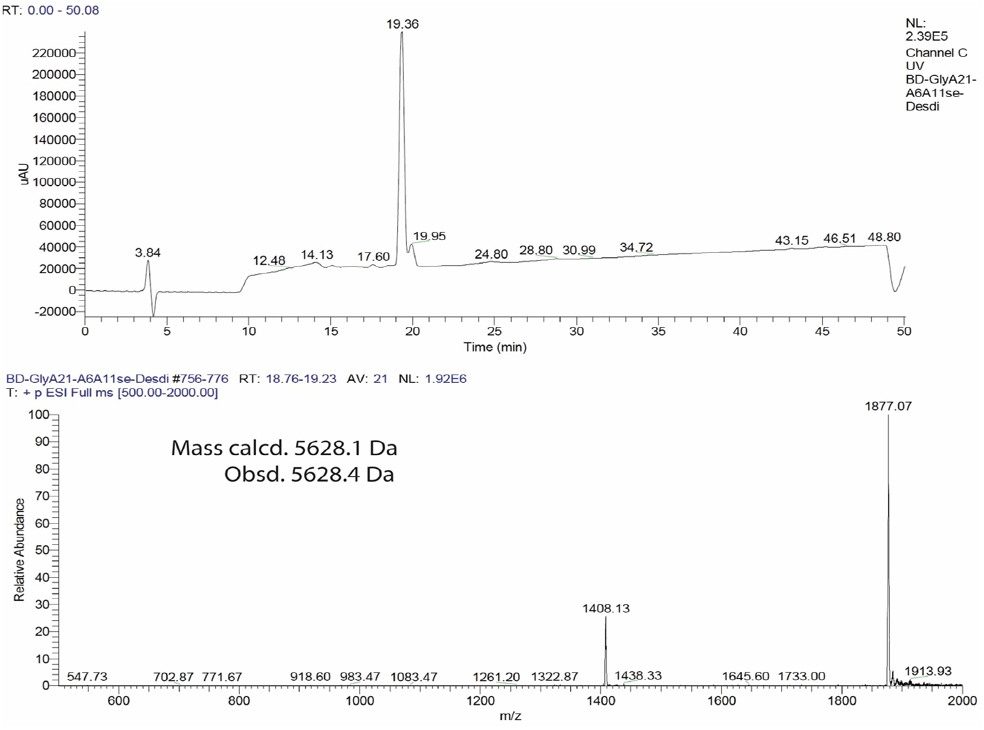
LC-MS analysis of folded 49-mer polypeptide [Se^A6^-Se^A11^]-Gly^A21^ Desdi SCI. HPLC chromatogram (monitored by UV absorbance at 215 nm) is shown at the top panel. ESI-MS spectra are shown in bottom panel.

**Figure 4.**
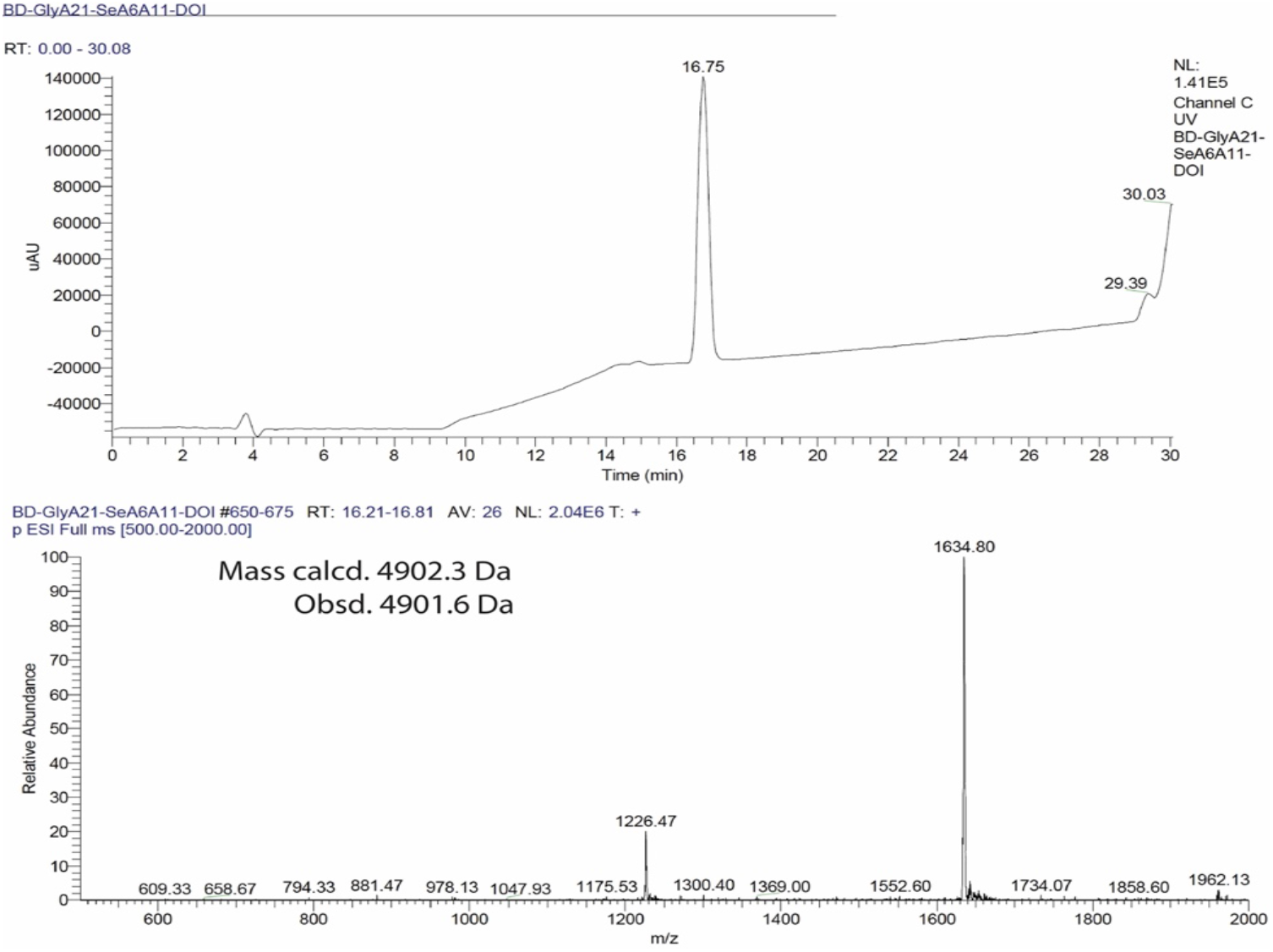
LC-MS analysis of [Se^A6^-Se^A11^]-Gly^A21^ desoctapeptide insulin (DOI). HPLC chromatogram (monitored by UV absorbance at 215 nm) is shown at the top panel. ESI-MS spectra are shown in bottom panel.

**Figure 5.**
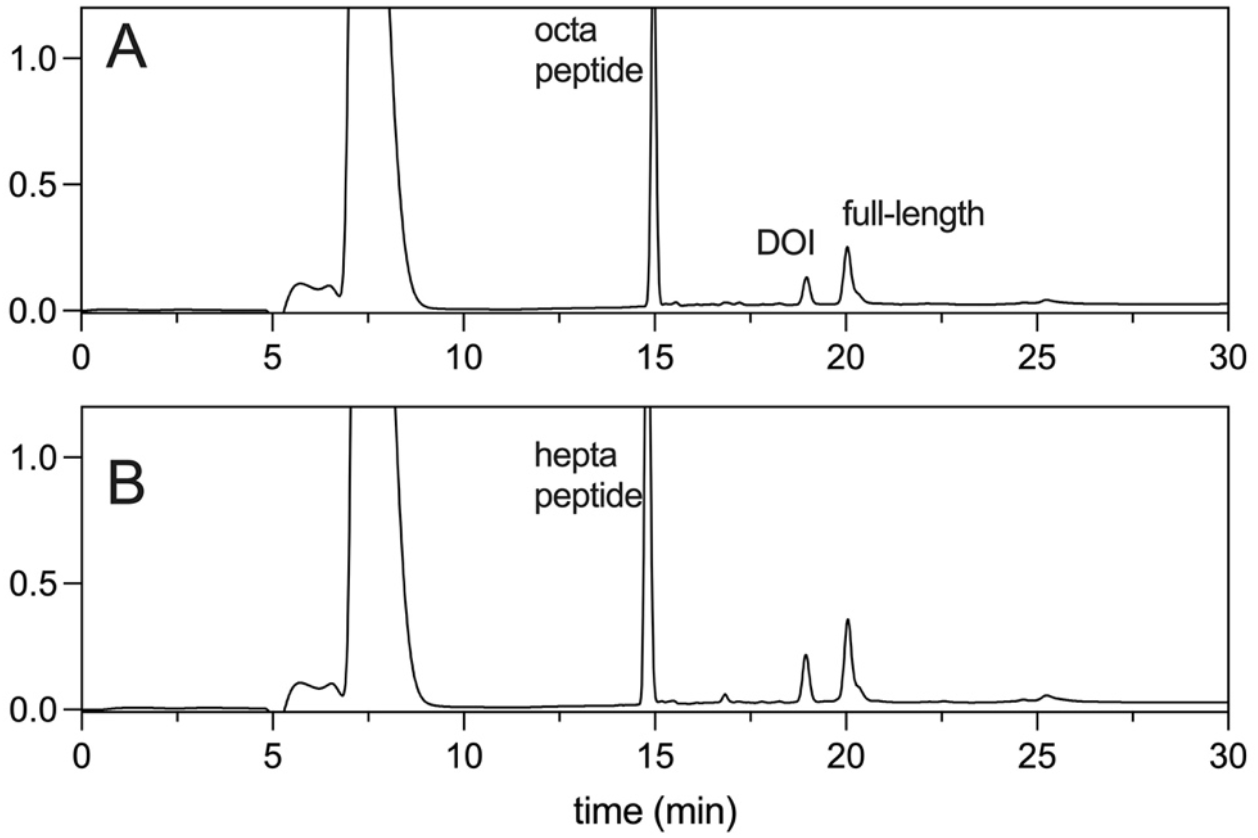
HPLC traces (monitored by UV absorbance at 215 nm) of semisynthesis. (A) [Se^A6^-Se^A11^]-Gly^A21^, Orn^B29^-insulin (Se-M1). (B) [Se^A6^-Se^A11^]-Gly^A21^, Orn^B29^, *des*-Thr^B30^ insulin (Se-M2).

**Figure 6.**
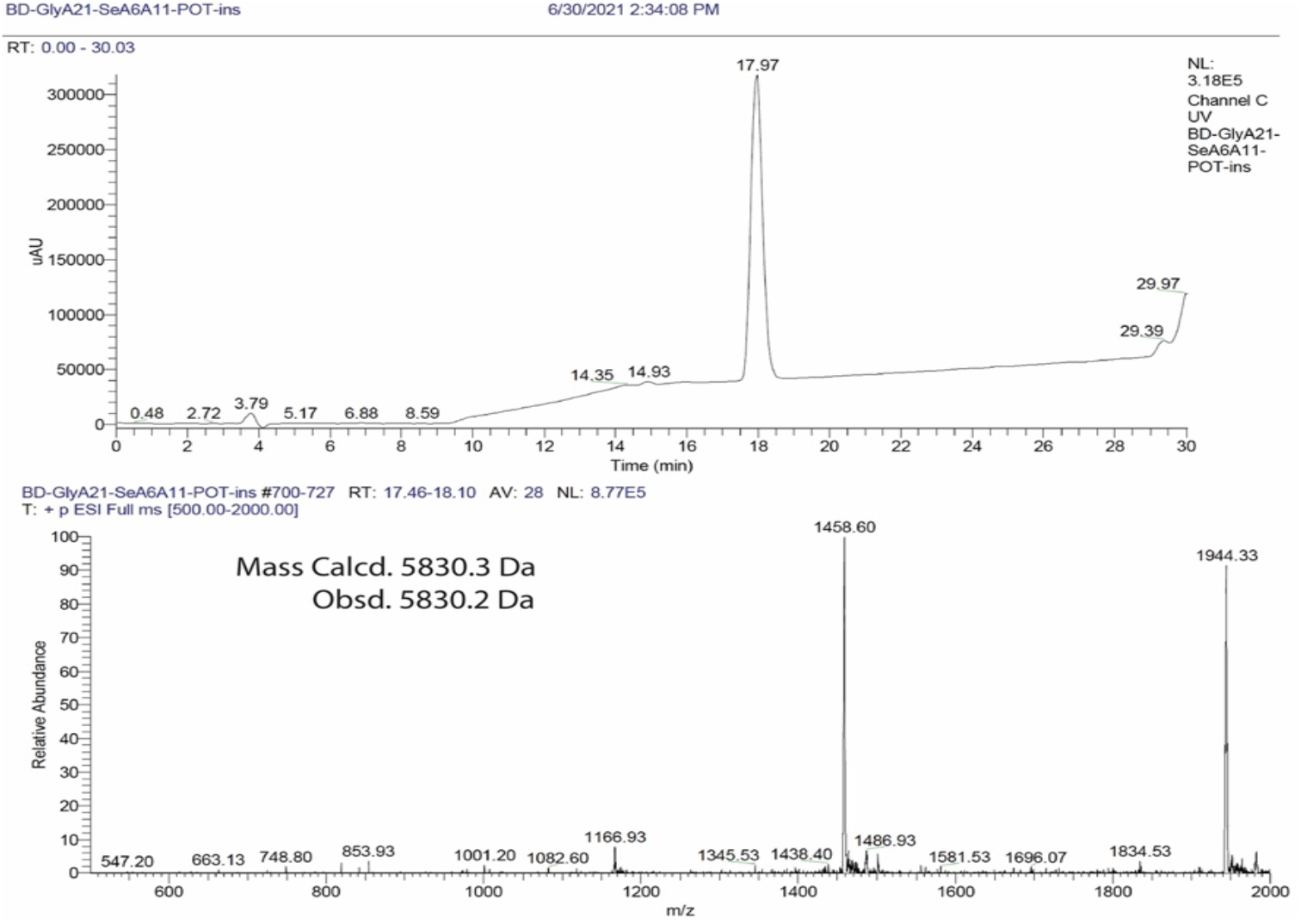
LC-MS analysis of [Se^A6^-Se^A11^]-Gly^A21^, Orn^B29^-insulin (Se-M1). HPLC chromatogram (monitored by UV absorbance at 215 nm) is shown at the top panel. ESI-MS spectra are shown in bottom panel.

**Figure 7.**
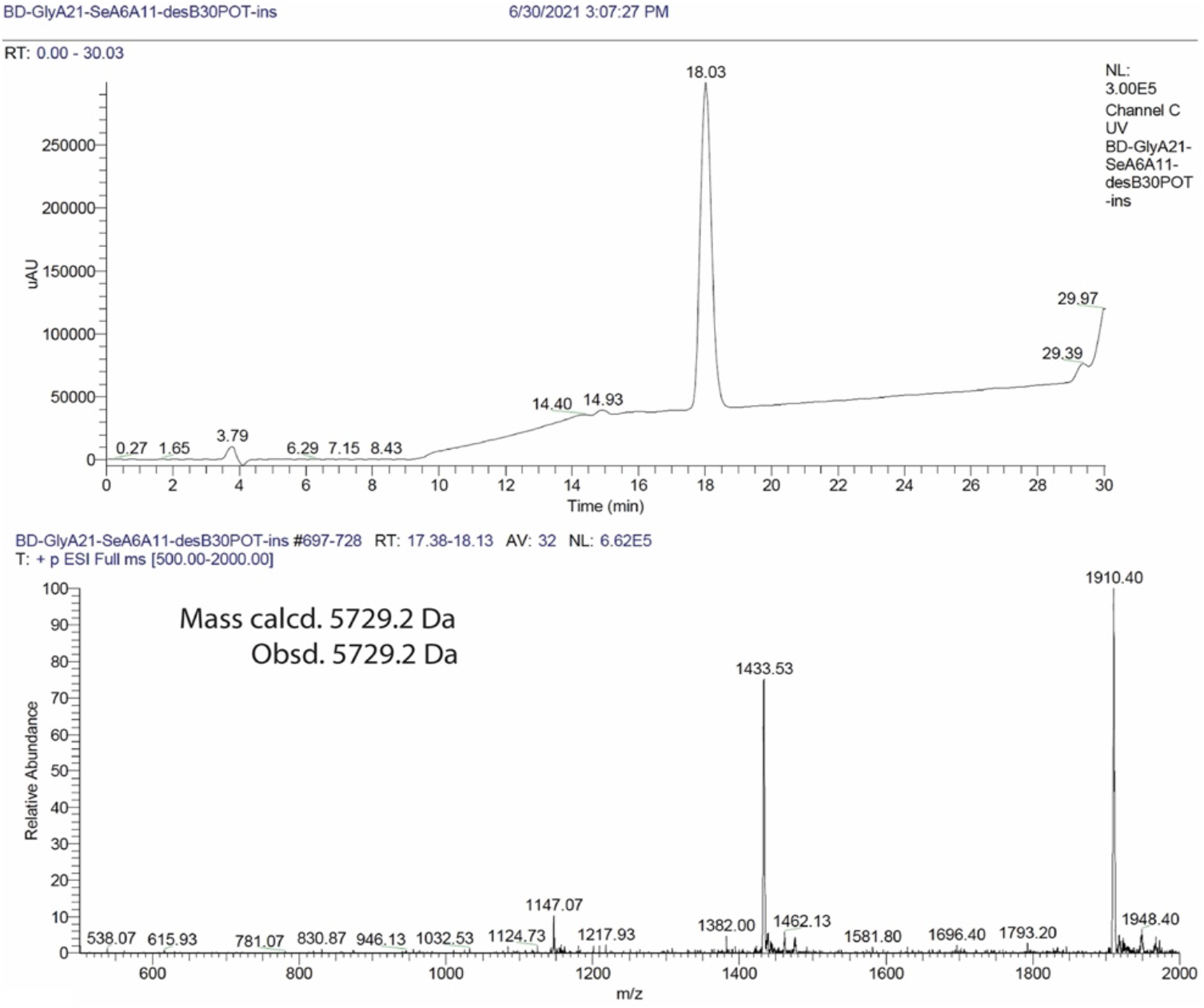
LC-MS analysis of [Se^A6^-Se^A11^]-Gly^A21^, Orn^B29^, *des*-Thr^B30^ insulin (Se-M2). HPLC chromatogram (monitored by UV absorbance at 215 nm) is shown at the top panel. ESI-MS spectra are shown in bottom panel.

### Se-M1 and Se-M2 exhibit native mitogenicities

As an assay of mitogenicity, cell signaling was probed by real-time quantitative reverse-transcriptase polymerase chain reaction (rt-qPCR) in two cell lines, rat L6 myoblasts overexpressing insulin receptor isoform A (L6 IR-A cells; 19) and human MCF-7 breast cancer cells (20); the latter express human IGF-1R and to a lesser extent IR isoforms A (IR-A) and B (IR-B) (20). Following exposure to 50 nM hormone (glargine, Se-glargine, Se-M1 and Se-M2 or control insulin analogs), transcriptional activation of mitogenic/proliferative pathways was assessed rt-qPCR measurements of cyclin mRNA abundances. Design of this assay was based on protein-level analyses of cyclins D1 and G2 (21) and subsequently adapted at the mRNA level (22).

A model insulin analog with native or baseline mitogenicity was provided by prandial analog insulin *lispro* (the active component of clinical formulation Humalog [Eli Lilly] (23)). Conversely, a model analog with enhanced mitogenicity was provided by Asp^B10^-insulin (also designated X10; 24), whose long-term administration in rats was found to lead to mammary carcinogenesis (24). Mitogenic signaling by Asp^B10^-insulin indeed led to a reduction in cyclin G2 and increase in cyclin D1 at the mRNA level (Fig. 8A,B). As expected, both glargine and Se-glargine displayed significant and similar mitogenicities (Fig. 8A,B). By contrast, WT insulin, insulin *lispro*, glargine derivatives M1, M2, Se-M1 and Se-M2 each exhibited baseline mitogenicities as indicated by comparable cyclin D1 activation and cyclin G2 repression.

**Figure 8.**
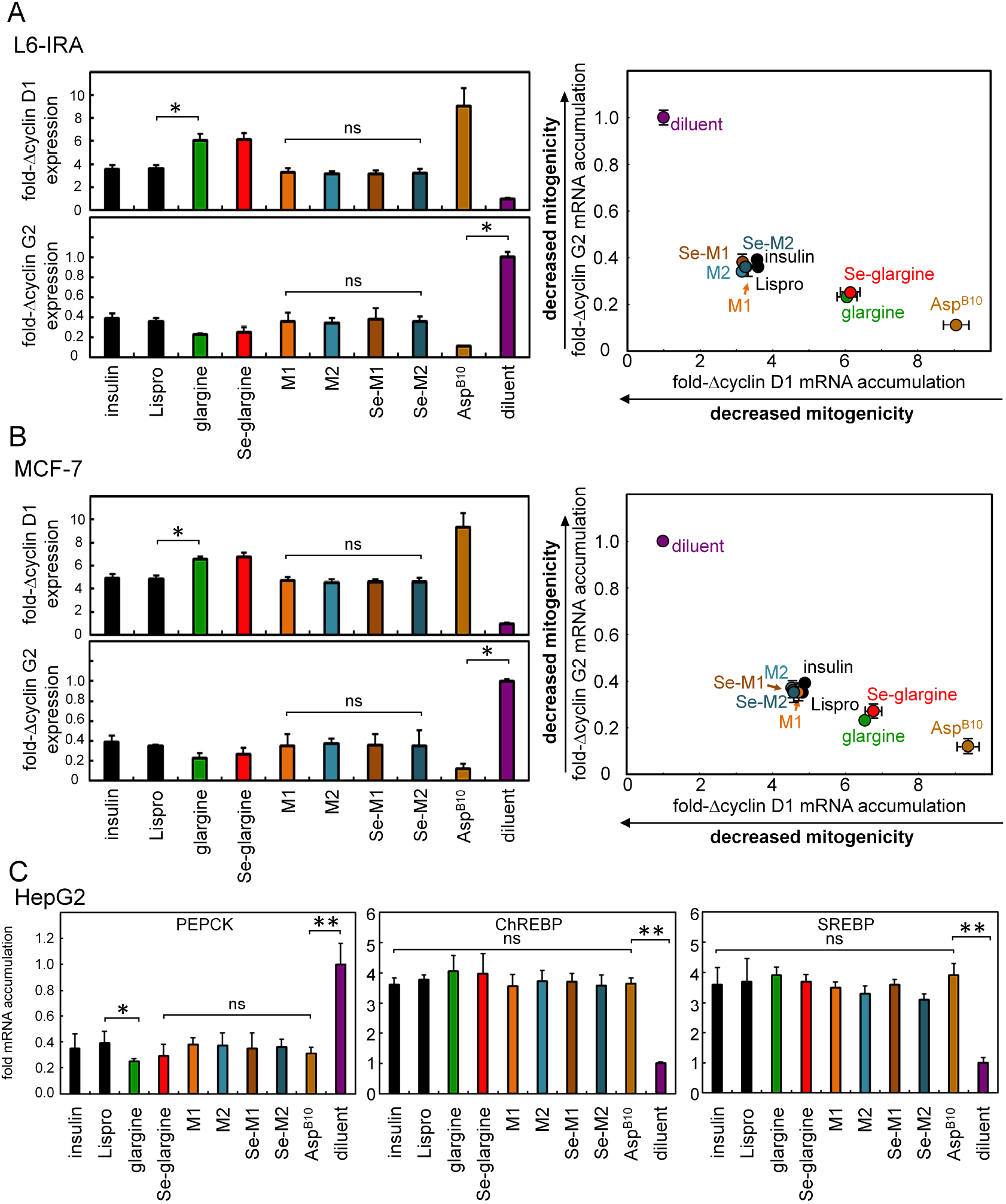
Cell biological assessment of insulin signaling. (A) The qPCR assay probed the cell proliferation markers, cyclin D1 and G2, to monitor the mitogenicity trends for insulin analogs. Human insulin and lispro serve as normal controls (black histograms) and analog Asp^B10^ provides the mitogenicity control for the assessment of hormone-induced IR-A signaling and activation of mitogenic pathways. L6 rat myoblasts stably expressing human IR isoform A (IR-A) (developed by De Meyts and co-workers (19)) and (B) MCF-7 human breast cancer cells expressing both IR isoforms (IR-A and IR-B) and high levels of IGF-1R were treated with analogs or controls. The increased accumulation of cyclin D1 and decreased cyclin G2 mRNA served as readouts for insulin-induced activation of cellular proliferation pathways. On the right side of each panels are assessments of insulin-driven cyclin D1 and G2 transcription in 2-D plots. Brackets designate *p* values: *, <0.05, or **, <0.01; ns indicates *p* values >0.05. (C)Transcriptional responses specific to PEPCK (*Left*) versus ChREBP and SREBP (*Middle* and *Right*, respectively). Decreased downstream mRNA accumulation of PEPCK and the increasing mRNA of ChREBP and SREBP reflect the activation stimulated by testing analogs in cellulare metabolism pathways. Asterisks (*) and (**) indicate *p* value <0.05 and <0.01. The “ns” indicates *p* value >0.05.

### Se-glargine retains native-like activity

Binding of insulin and insulin analogs to the insulin receptor (IR) on the surface of cells leads to receptor autophosphorylation (for review, see ref 25,26). This reaction reflects the hormone-dependent activation of the intracellular receptor tyrosine kinase (26), the first step in signal propagation from outside of the cell to inside. Dose-dependent measurement of the ratio of phosphor-IR (pIR) to total IR thus provides a quantitative measurement of relative hormone activity and is readily implemented as 96-well plate assay (Fig. 9A). Dose-response relationships of insulin *lispro* (KP) and native glargine were similar; any small differences were not statistically significantly different (Fig. 9B). Similar results were obtained in assays of Se-glargine vs. Se-M1 (Fig. 9C) and insulin *lispro* vs. Se-M2 (Fig. 9D), providing evidence for their native activities.

**Figure 9.**
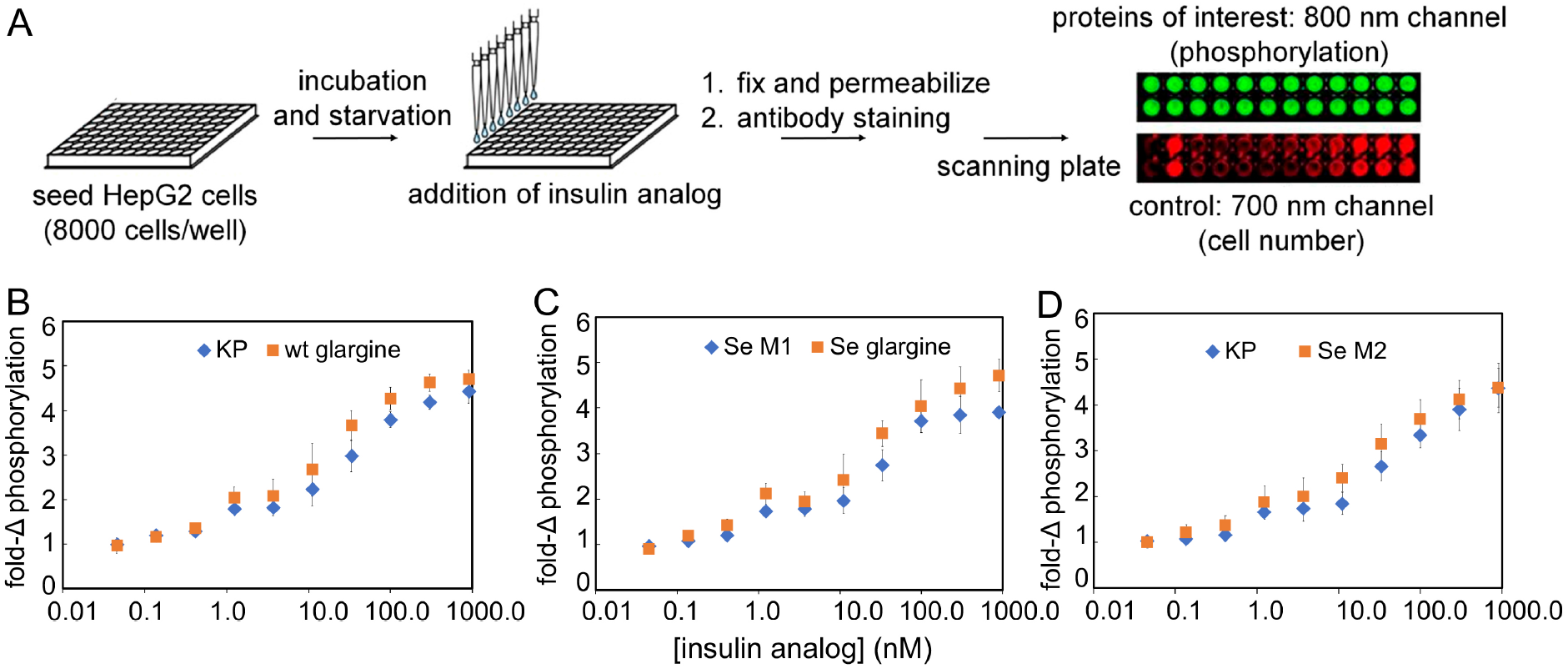
Insulin signaling and dose–response relationships. (A) Schematic outline of in-cell western blotting assay probing IR autophosphorylation stimulated by insulin analogs. Flowchart illustrates procedure to assess hormone-induced phosphorylated IR via 800 nm readouts and the assay applied cell number control using 700 nm readouts. (B-D) Respectively show similarity in activities for various insulin analogs: (B) insulin *lispro* (KP; *blue diamond*) versus native glargine (wt glargine; *red square*), (C) Se-M1 (*blue diamond*) versus Se-modified glargine (Se glargine; *red square*), and (D) KP (*blue diamond*) versus Se-M2 (*red square*). In each pair-wise comparisons, *p*-values were >0.05 and thus not significant. Ligand concentrations are shown in horizontal axis. Ligand doses: 0.05, 0.14, 0.41, 1.2, 3.7, 11.1, 33.3, 100, 300, and 900 nM; assays were performed in triplicate.

### Insulin-Mediated Metabolic Gene Regulation

Insulin signaling in hepatocytes extends to metabolic transcriptional regulation (phosphoenolpyruvate carboxykinase; PEPCK, carbohydrate-response-element binding protein; ChREBP, and sterol-response-element binding protein; SREBP) (for review, see ref 27). This signaling network can be recapitulated in HepG2 cells (Fig. 8C) (28,29). Under hypoglycemic conditions, for example, HepG2 cells exhibit exogenous insulin-specific suppression of gluconeogenesis-related gene expression as probed here by *PEPCK* gene expression (29): insulin signaling thus overrides hypoglycemic activation of the *PEPCK* gene in these cells. Such suppression was broadly observed among the present set of insulin analogs (at left in Fig. 8C). Similarly, under normoglycemic conditions, insulin signaling activates transcription of genes encoding ChREBP and SREBP, mediators of the hormonal control of lipid biosynthesis (29). No significant differences were observed among analogs (middle and right in Fig. 8C).

## Discussion

Selenocysteine modified at internal cysteine residues (A6 and A11) in human insulin was previously found to exhibit increased thermodynamic stability at neutral pH relative to WT insulin (ΔΔG_u_ *ca*. 1 kcal/mol) (9). In the present study we have extended this finding to insulin glargine under acidic conditions. This basal insulin analog is formulated at pH 4.0 and undergoes isoelectric precipitation at the neutral pH of the subcutaneous space to provide a long-lived depot of insulin (7). This basal formulation is in broad clinical use in both the developed and developing worlds (5,6). We sought to engineer enhanced stability (both dynamic and thermodynamic) because at pH 4.0 insulin glargine is not protected by zinc-hexamer assembly from physical and chemical degradation: protonation of His^B10^ precludes its coordination of zinc cations (Zn^2+^) (30).

The present studies of ^1^H-^2^H amide proton exchange in D_2_O provide evidence of a smaller but significant increase in thermodynamic stability (as probed by global ^1^H-^2^H amide proton exchange at position B18; red in Table 1) and damping of subglobal conformational fluctuations in the central B-chain α-helix and C-terminal A-chain α-helix. Both effects can delay degradation of an insulin formulation (31,32). Because of the exponential function, even differences of 0.2-0.4 kcal/mole would be predicted to at least double the shelf life of an insulin formulation at room temperature. It would be of future interest to repeat these studies at 40 °C wherein the protective effects of the diselenide bridge may be enhanced relative to 25 °C. Such an effect would be of interest in relation to global access of patients to insulin, as transient exposure of insulin products to high temperatures shortens its shelf life even at or below room temperature (31,32). The importance of this issue is magnified by an emerging global pandemic of obesity-related Type 2 diabetes mellitus (33,34).

The present ^1^H-^2^H amide-proton exchange studies may underestimate the degree of protection afforded by the A6-A11 diselenide bridge, as conformational fluctuations in the A1-A10, B1-B8 and B20-B30 were not probed—and yet may open the protein molecule to chemical or physical degradation. In the future it would be of interest to extend these studies to rapid sites of amide-proton exchange, such as through tandem mass-spectrometric methods to monitor such exchange (35). It would likewise be of interest to measure the degradation rates of native glargine versus Se-glargine in a clinical formulation under real-world conditions (36).

Diselenide bridges are intrinsically more resistant to reduction than disulfide bridges (9). Redox chemistry is nonetheless tangential to the present application of selenium chemistry. Indeed, substitution of an external or solvent-exposed disulfide bridge (cystine A7-B7) by the corresponding diselenide bridge was found not to stabilize human or bovine insulin (37). We therefore ascribe the stabilizing effect of the A6-A11 diselenide bridge to its more efficient core packing. This finding exploits suboptimal packing of native insulin monomer near cystine A6-A11 (Fig. 10) (30). Since selenium has a slightly larger atomic radii and longer Se-Se bond length (38), the modified bridge would be expected to better fill the native microcavities, thereby mitigating baseline cavity penalties (39). We predict that this approach will generally apply to all classes of insulin analogs.

**Figure 10.**
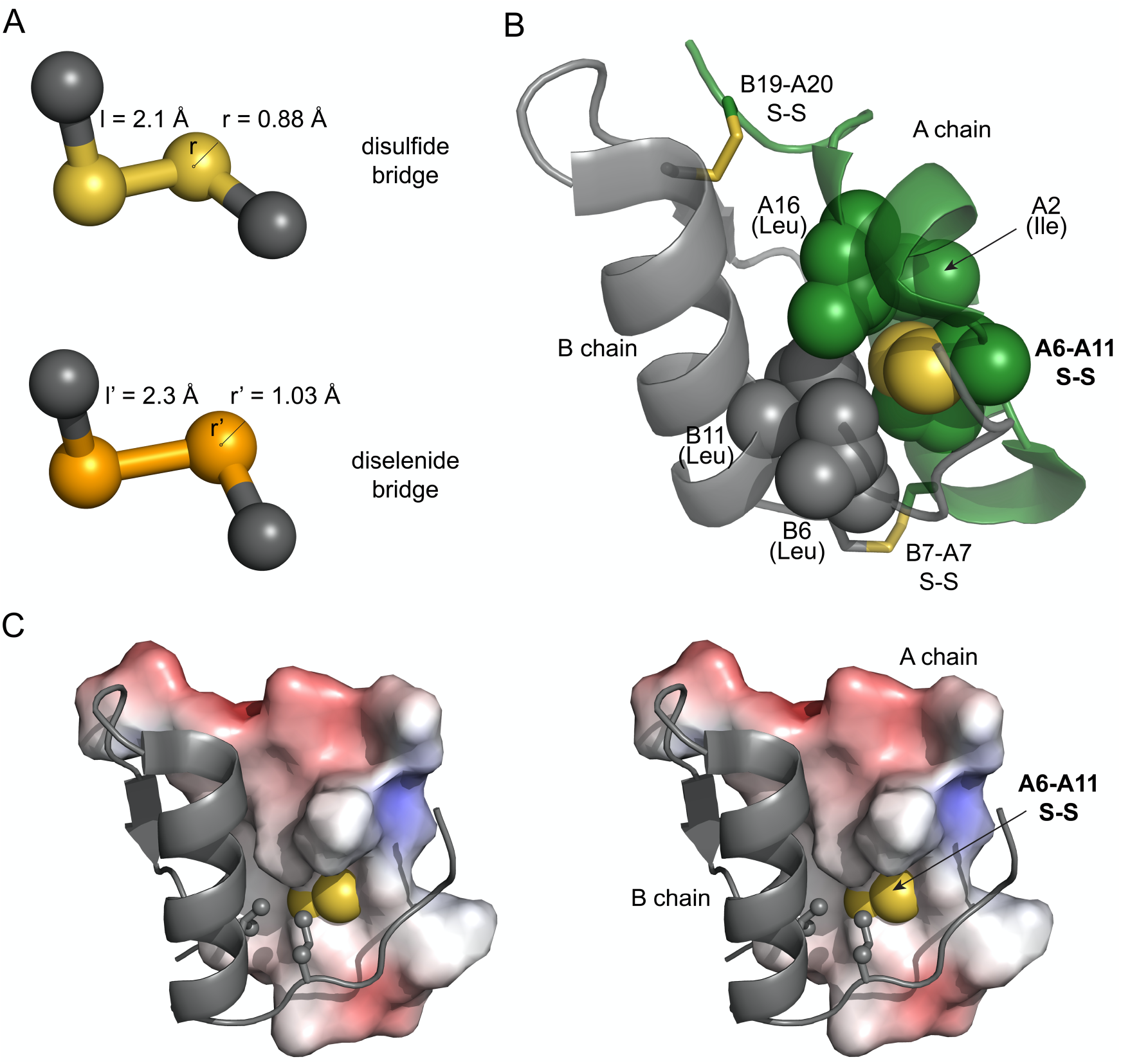
Diselenide protein engineering to optimize the insulin structure. (A) comparison of atomic radii and bond lengths in disulfide and diselenide bridges. Selenium has relatively larger atomic radius and its longer bond length in diselenides makes them ideal candidate for protein engineering. (B) Cartoon struture showing the cavity near Cys^A6^-Cys^A11^ disulfide in T-state of insulin monomer (PDB 4INS). A chain (*green*), B chain (*gray*), disulfides A7-B7 and A20-B19 shown as sticks. Residues Cys^A6^-Cys^A11^ and neighboring residues Leu^B6^, Leu^B11^, Ile^A2^ and Leu^A16^ are shown as spheres. Sulfur atom is shown in *yellow* color. (C) Stereoview model showing environment around Cys^A6^ and Cys^A11^. B chain shown in gray as cartoon, whereas A chain is shown as electrostatic surface (calculated in the absence of Cys^A6^ and Cys^A11^ side chain). Side chains of Leu^B6^ and Leu^B11^ are shown as sticks with the methyl groups in sphere representation with one third of its van der Waals radii.

### Experimental Procedures

#### Materials and Methods

Fmoc amino acids were purchased from Gyros Protein Technologies. Protecting groups used were: Arg(Pbf), Asn(Trt), Cys(Trt), Gln(Trt), Glu(OtBu), His(Trt), Lys(Boc), Ser(tBu), Thr(tBu), Tyr(tBu), Orn(Boc). Fmoc-Sec(Mob)-OH was purchased from Chem-Impex International. Fmoc-Gly-wang resin (0.68 mmol/g loading), 6-chloro-1-hydroxybenzotriazole (6-Cl-HOBt) and *N,N’*-diisopropylcarbodiimide (DIC) were obtained from Gyros protein Technologies. 2-chlorotritylchloride resin (1.12 mmol/g loading), piperidine and trifluoroacetic acid were purchased from Chem-Impex International. Diethyl ether, dichloromethane (DCM), *N,N*-dimethylformamide (DMF), acetonitrile (HPLC-grade), and guanidine hydrochloride were purchased from Fisher. Trypsin, TPCK treated was purchased from Thermo Scientific. All other reagents were purchased from Sigma-Aldrich and were of the purest grade available.

#### Analytical and semipreparative HPLC conditions

Analytical reversed-phase HPLC was performed using a Waters 1525 Binary HPLC system. All chromatographic separations were performed on a C8 Proto (4.6x250 mm) 300 Å, 5 μm, Higgins Analytical Inc. column, using 25-50% Solvent B in Solvent A over 35 minutes. (solvent A = 0.1% TFA in water, solvent B = 0.1% TFA in acetonitrile) at a flow rate of 1.0 mL/min with detection by UV absorption at 215 nm. For semipreparative HPLC following conditions were used: TARGA C8 5μm (250×10mm) Higgins Analytical, Inc. column at flow rate of 4 mL/min.

#### Preparative HPLC conditions

Waters 2545 Quaternary pumping system equipped with FlexInject was used for the preparative HPLC purification. Chromatographic separations were performed on a C4 Proto (20×250 mm) 300 Å, 10 μm, Higgins Analytical Inc. column, using 25-50% Solvent B in Solvent A over 35 minutes at a flow rate of 20 mL/min with detection by UV absorption at 215 nm. Fractions containing the desired products were identified by analytical LC and mass spectrometry, then combined and lyophilized.

#### LC-MS

LC-MS analysis was performed on LCQ Advantage Ion Trap Mass Spectrometer System coupled with Agilent 1100 Series HPLC system. Masses were obtained by online electrospray mass spectrometry. MS data shown were collected across the entire principal UV absorbing peak in each chromatogram.

### Peptide synthesis for the preparation of Se-glargine M1 and M2

#### DesDi Sequence

##### FVNQHLCGSHLVEALYLVCGERGFFYTK-GIVEQUCTSIUSLYQLENYCG

[Se^A6^-Se^A11^]-Gly^A21^ DesDi-single chain insulin was synthesized on Tribute (Gyros Protein technologies) peptide synthesizer. All the amino acids were 10x excess relative to the resin except the precious Fmoc-Sec(Mob)-OH which was used in 3-fold excess. 6-Cl-HOBt and DIC (1:1, equimolar with respect to Fmoc-protected amino acids) were used as the coupling agents and 20% piperidine in DMF used for deprotection. At the end of the synthesis, peptides were cleaved with TFA cocktail 2.5% *v/v* of 2,2′-(ethylenedioxy)diethanethiol, triisopropylsilane, anisole, water and 2,2′-dithiobis(5-nitropyridine) (DTNP, 2 equivalents per selenocysteine; ref 40). Cleavage mixture was precipitated with ether (5 fold with respect to TFA) and solid was isolated by centrifugation. Precipitate was further washed with twice with ether and dried in vacuo.

Octapeptide (sequence: GFFYTPOT) and heptapeptide (sequence: GFFYTPO) (O = Orn) were synthesized on 2-chlorotritylchloride resin. After the synthesis and final N^α^-Fmoc deprotection, the resin was cleaved using TFA cocktail and worked-up as described above. The resulting peptides were dissolved in 1:1 solvents A/B and lyophilized. This was used in semi-synthesis without further purification.

### Folding of DesDi-precursor and DOI-generation

#### DesDi

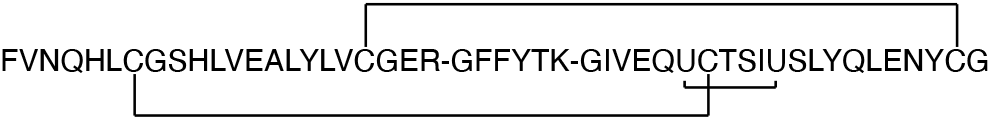

#### DOI

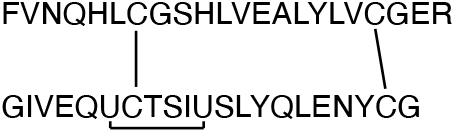

The crude 49-mer peptide, after the Fmoc-SPPS, was subjected to folding conditions: 0.1 mM peptide, 20 mM glycine, 2 mM cysteine at pH 10.5 for 16 hr (18). After HPLC indicated the sharp early eluting peak, the reaction mixture acidified to pH 3.0, filtered (0.22 μ) and purified on a reverse phase (RP) C4-preparative column. To generate the *des*-octapeptide insulin (DOI), the single chain DesDi precursor was treated with trypsin-TPCK (20% *w/w*) in a buffer containing 1 M urea and 0.1 M ammonium bicarbonate for 24 hr at room temperature. After HPLC indicated the completion of the cleavage reaction, DOI was purified using RP-HPLC.

### Semi-synthesis of Se-glargine M1 and M2

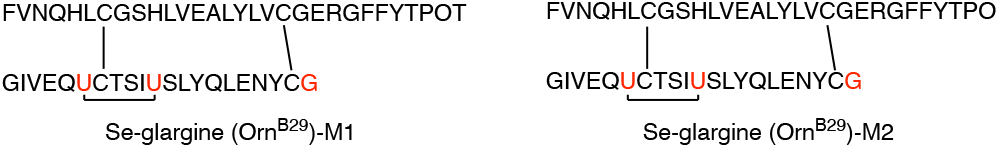

The DOI from the above step was combined with octapeptide GFFYTPOT (for M1) or heptapeptide GFFYTPO (for M2) in presence of 10% *w/w* trypsin-TPCK as reported previously (16). Orn^B29^ was used in place of Lys to enable the semi-synthesis preparation of these analogs. After the HPLC indicated the conversion to more than 50-60%, the reaction diluted in 0.1% TFA, 6 M guanidine hydrochloride (GuHCl) and purified by semi-preparative HPLC.

### ^1^H-^2^H Exchange NMR Assay in D_2_O

^1^H-^2^H exchange was performed in 100% D_2_O in 10mM deuterated acetic acid at pH 3.0 (direct meter reading) at 25°C. Lyophilized samples of native glargine or Se-glargine samples (lyophilized from the samples in H_2_O in 10 mM acetic acid) were placed on ice followed by addition of chilled D_2_O containing 10 mM deuterated acetic acid. A series of 1D proton and 2D TOCSY spectra were collected to measure exchange rate of amide protons as a function of peak intensity; 1D ^1^H spectra were collected at 2 minutes intervals to assess very fast exchange of amide protons, while 2D TOCSY spectra were acquired to assess amide protons with slower exchange rates. NMR data were processed using Topspin 4.0.6 (Bruker Biospin) and analyzed with Sparky software (41). Protection factors were calculated from the ratio of measured exchange rate and intrinsic exchange rate. Estimates of protein stability were derived from the relationship between PF and exchange free energy (ΔG_HX_) (42,43).

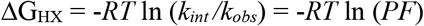

where R is the Rydberg constant, T is absolute temperature, *k*_*int*_ *i*s intrinsic rate constant at a given residue, and *k*_*obs*_ is the observed ^1^H-^2^H -exchange rate constant.

### Cell culture

Three mammalian cell lines from human (HepG2 and MCF-7) and rat (L6) were applied in the cell biology experiments. Human hepatocellular carcinoma cell line HepG2 was cultured in Dulbecco’s Modified Eagle Medium (DMEM), supplemented with 10% fetal bovine serum (FBS), 1% penicillin/streptomycin as recommended by the American Type Culture Collection (ATCC). A protocol employing 24-h serum starvation was applied from (29,43) except that FBS was applied at 70-75% confluence. After starvation, cells were treated in parallel with a set of insulin analogs in serum-free medium.

Signaling activities of cell proliferation responses stimulated by the analogs were tested in the human breast adenocarcinoma cell line, MCF-7. MCF-7was cultured in EMEM medium, supplemented with 10% FBS, 1% penicillin/streptomycin, and sodium pyruvate (1 mM) but without insulin. 24-hours serum-starving protocol using the culture medium except FBS was applied after reaching 70-75% confluence (approximately 6X10^6^ cells per 10-cm dish). After the starving, cells were treated with the serum-free medium containing 50 nM of tested insulin analogs simultaneously. For mitogenicity qPCR assays, the treatment time was 8 hours. After the remove of analog-containing medium, the qPCR lysis kit (Bio-Rad) was applied to prepare the cell lysate for one-step qPCR analysis.

Cell proliferation responses was also tested in L6 (rat myoblast cell line) with exogenous human IR expression (L6-IRA). L6-IRA cells will be cultured in DMEM medium supplemented with 10% FBS, and G418 providing selection condition. 24-hours serum-starving protocol using culture medium except FBS will be applied after reaching 70-75% confluence (approximately 0.8X10^6^ cells per well). After the starving, serum-free medium containing 50nM of tested insulin analogs will be added in each well simultaneously. Time of treatment is 8 hours and insulin-containing medium will be removed. The cell-lysis protocol using qPCR lysis kit (Bio-Rad) was applied for preparing for one-step qPCR analysis.

### Real-time qPCR Assays

Following serum starvation, HepG2 cells were treated with medium containing an insulin analog (50 nM) for 8 h. In studies related to possible glucose responsiveness and lipid metabolism, the cells were treated with analogs for 3 or 4 hours in media containing either low or normal glucose concentrations. Readouts were provided by downregulation of PEPCK and G6P and upregulation of ChREBP and SREBP. mRNA (messenger ribonucleic acid) abundances were measured in triplicate by quantitative polymerase chain reaction (qPCR). Samples were prepared as described by the vendor (One-Step rt-PCR reagent kits; Bio-Rad).

The following sets of primers were used: (PEPCK), GTTCAATGCCAGGTTCCCAG and TTGCAGGCCAGTTGTTGAC; (ChREBP), AGAGACAAGA-TCCGCCTGAA and CTTCCAGTAGTTCCCT-CCA; (SREBP), CGACATCGAAGACATGCTTCAG and GGAAG-GCTTCAAGAGAGGAGC; and (GAPDH), ATGGTTTACATGTTCCAATAT and ATGAGGTCCACCACCCTGGTTG.

Mitogenesis probing gene expressions in MCF-7 cell line stimulated by insulin analogs was under 50nm of analog working concentration with 8 hours of treatment. The transcription was measured in triplicate by quantitative real-time-Q-rtPCR (qPCR). Sample preparation were followed the instruction of One-Step rt-PCR reagent kits as described by the vendor (Bio-Rad).

The following sets of primers were used: (Cyclin D1), AATGACCCCGCACGATTTC and TCAGGTTCAGGCCTTGCAC; (Cyclin G2), ATCGTTTCAAGGCGCACAG and CAACCCCCCTCAGGTATCG; (GAPDH): AGCCGAGCCACATCGCT and TGGCAACAATATCCACTTTACCAGAGT; (TFIID), GCACAGGAGCCAAGAGTGAA and TCACAGCTCCCCACCATGTT.

Mitogenesis probing gene expressions in L6-IRA cell line stimulated by insulin analogs was under 50 nm of analog working concentration with 8 hours of treatment. The transcription was measured in triplicate by quantitative real-time-Q-rtPCR (qPCR). Sample preparation were followed the instruction of One-Step rt-qPCR reagent kits as described by the vendor (Bio-Rad).

The following sets of primers were used. (Cyclin D1), GCCGAGTGGAAACTTTTGTCG and CGGGAAGCGTGTACTTATCCT; (Cyclin G2), GCAAGAAAAGAAGCCAAGCT and TGACCAAGAGGCAAAATAAAATCAA; (GAPDH), GACATGCCGCCTGGAGAA and GCCCAGGATGCCCTTTAGT; (TFIID), CTGAGGGGGCAATGTCTAAC and GGGCAGCTAGTGAGATGAGC.

### Plate-based Fluorescence Immunoblotting Assay

The assay was designed to assess hormone-induced IR autophosphorylation via in-cell illumination readouts. Human liver-cancer derived HepG2 cells were seeded (∼8000 cells per well) into a 96-well assay black plate with clear bottom, and grown in tissue culture (Fisher). After starving in 100 μl of plain Hank’s Balanced Salt Solution (HBSS) for two hours at 37 °C, 100 μl of a given insulin analog at a specific dose (in the range 0.5-100 nM) were applied to each well and incubated for 20 minutes at 37 °C. After removing the insulin solution, 150 μl of 3.7% formaldehyde (Fisher) was added to each well, and plates were incubated for 20 minutes at 37 °C; 200 μl of 0.1 % Triton-X-100 detergent solution (Sigma) was then added to mediate cell permeabilization under same incubation conditions. 100-μl Odyssey Blocking Buffer (LI-COR) was added to fixed cells after permeabilization. The blocking procedure was implemented for 1 hour at room temperature on gentle agitation in an orbital shaker.

After blocking, the fixed cells were incubated with the primary antibody (10 μl anti-pTyr 4G10 into 20 mL Blocking Buffer) overnight at 4°C. After a wash, the secondary antibody (anti-mouse-IgG-800-CW antibody (Sigma) into 25 mL Blocking Buffer) was added. This enabled detection at 800 nm (red emission frequency) to probe the extent of pTyr modification. DRAQ5 (Fisher) was also applied to enable 700 nm emission (green) as a control to estimate cell number. These respective fluorescence signals were determined at 700 and 800 nm using an LI-COR Infrared Imaging system (Odyssey) with offset set to 4 mm and intensity set to “auto”.

## Abbreviations

Amino acids are designated by standard three-letter or one-letter code, with selenocysteine abbreviated as Sec or U. Residues within insulin are typically indicated by three letter code with chain and residue number in superscript; *e*.*g*., asparagine or glycine at position A21 of the A chain would be designated Asn^A21^ or Gly^A21^.

## Author Contributions

Protein chemical synthesis, purifications were performed by B.D. and O.W.-K. with the advice of M.A.W. and N.M. Cell-biological assays were performed by Y-S.C with advice from M.A.W. NMR studies were performed by Y.Y and M.A.W. The overall program of research was guided by N.M. and M.A.W. All authors contributed to preparation of the manuscript.

## Acknowledgements

We thank members of our respective laboratories for discussion. O.W-K was supported by the Kaete Klausner Ph.D. scholarship. N.M. acknowledges the financial support of the Israel Science Foundation (1388/22); M.A.W. acknowledges the support of the U.S. National Institutes of Health (NIH R01 DK04949).

## FOOTNOTES

*Disclosure*. O.W.-K., N.M. and M.A.W. hold a US patent number PCT29920-315514, entitled “Stabilization of prandial or basal insulin analogs by an internal diselenide bridge,” which is supported by this study.

